# Genome evolution in bacteria isolated from million-year-old subseafloor sediment

**DOI:** 10.1101/2020.12.19.423498

**Authors:** William D. Orsi, Tobias Magritsch, Sergio Vargas, Ömer K. Coskun, Aurele Vuillemin, Sebastian Höhna, Gert Wörheide, Steven D’Hondt, B. Jesse Shapiro, Paul Carini

**Author notes:** **Corresponding authors:** Prof. Dr. William D. Orsi, Ludwig-Maximilians-Universität München, Department of Earth and Environmental Sciences, Paleontology & Geobiology, Richard-Wagner-Strasse 10, 80333 Munich, Germany. Dr. Paul Carini, Address: University of Arizona, Department of Environmental Science, School of Plant Science, BIO5, Institute. Tucson, Arizona 85721, E-Mail.

## Abstract

Beneath the seafloor, microbial life subsists in isolation from the surface world under persistent energy limitation. The nature and extent of genomic evolution in subseafloor microbes has been unknown. Here we show that the genomes of *Thalassospira* bacterial populations cultured from million-year-old subseafloor sediments evolve by point mutation, with a relatively low rate of homologous recombination and a high frequency of pseudogenes. Ratios of synonymous to non-synonymous mutation rates correlate with the accumulation of pseudogenes, consistent with a dominant role for genetic drift in the subseafloor strains, but not in type strains of *Thalassospira* isolated from the surface world. Our findings demonstrate that the long term physical isolation of these bacteria, in the absence of recombination, has resulted in clonal populations that evolve consistent with ‘Mullers Ratchet’, whereby reduced access to novel genetic material from neighbors has resulted in fixation of new mutations that accumulate in genomes over millions of years.

**Significance statement:** The nature and extent of genomic evolution in subseafloor microbial populations subsisting for millions of years below the seafloor is unknown. Subseafloor populations have ultra-slow metabolic rates that are hypothesized to restrict reproduction and, consequently, the spread of new traits. Our findings demonstrate that genomes of cultivated bacterial strains from the genus *Thalassospira* isolated from million-year-old abyssal sediment exhibit greatly reduced levels of homologous recombination, elevated numbers of pseudogenes, and genome-wide evidence of relaxed purifying selection. These substitutions and pseudogenes are fixed into the population, suggesting the genome evolution of these bacteria has been dominated by genetic drift, whereby under long-term physical isolation in small population sizes, and in the absence of homologous recombination, newly acquired mutations accumulate in the genomes of clonal populations over millions of years.

## Main text

The subseafloor biome contains a large fraction of all prokaryotic cells on Earth totaling circa 10^29^ cells (*1*). This biome subsists over geological timescales under persistent energy limitation (*2–6*). Whether evolution and ecological differentiation occurs in microbial populations below the seafloor has remained controversial. It is generally agreed that extreme energy limitation restricts metabolic activity and growth (*2–6*), which are necessary for new mutations to propagate through populations to foster ecological differentiation and speciation (*7*). However, there has been very little direct examination of this issue at the low metabolic rates and long timescales characteristic of subseafloor life (*8*). Experimental evidence exists for bacterial evolution under energy limitation on laboratory timescales (*9, 10*), but a recent metagenomic analysis showed that energy limitation and reduced growth restricted the spread of new mutations through microbial communities over 5,000 years in the upper 2 meters of anoxic sediment from Aarhus Bay (Denmark) (*11*). There have been no direct studies of mutation, homologous recombination, and evolution in microbial communities of the deeper and older sediment that dominates the subseafloor. Here, we used the genomes of bacteria isolated from million-year-old subseafloor abyssal clay sediments to investigate the nature of genome evolution in subseafloor bacteria that persist under extreme energy limitation over long timescales.

Newly acquired mutations of functional significance can sweep through relatively fast-growing bacterial populations in surface world habitats and influence ecological differentiation (*12*). However, it is unclear whether such sweeps occur in ancient subseafloor sediment given the comparably slow subseafloor bacterial biomass turnover rates that are estimated to be on thousand-year timescales (*13*). The metabolic rates of microbes persisting in deep-sea abyssal clay are amongst the lowest observed in the subseafloor biosphere, such that these sediments are often oxic through the entire sediment column to the underlying oceanic crust and the microbes live near the low-energy limit to life (*3, 4*). We cored a 15 m sedimentary sequence of oxygenated abyssal clay at a water depth of 6,000 m in the North Atlantic where the average sedimentation rate is an estimated 1 meter per million years (*14*). Thus, the deepest sediment sampled was deposited ca. 15 million years ago (mya). The relatively slow drawdown of O_2_ with increasing depth at this site (Fig. 1A) primarily reflects oxidation of organic matter by aerobic microbes. Abyssal clay is characterized by very low permeability and extremely small pore diameter (*15*), which physically isolates the subseafloor microbial communities from the surface and the microbes within it from each other (see ‘sediment physical properties’ in the SI).

**Figure 1:**
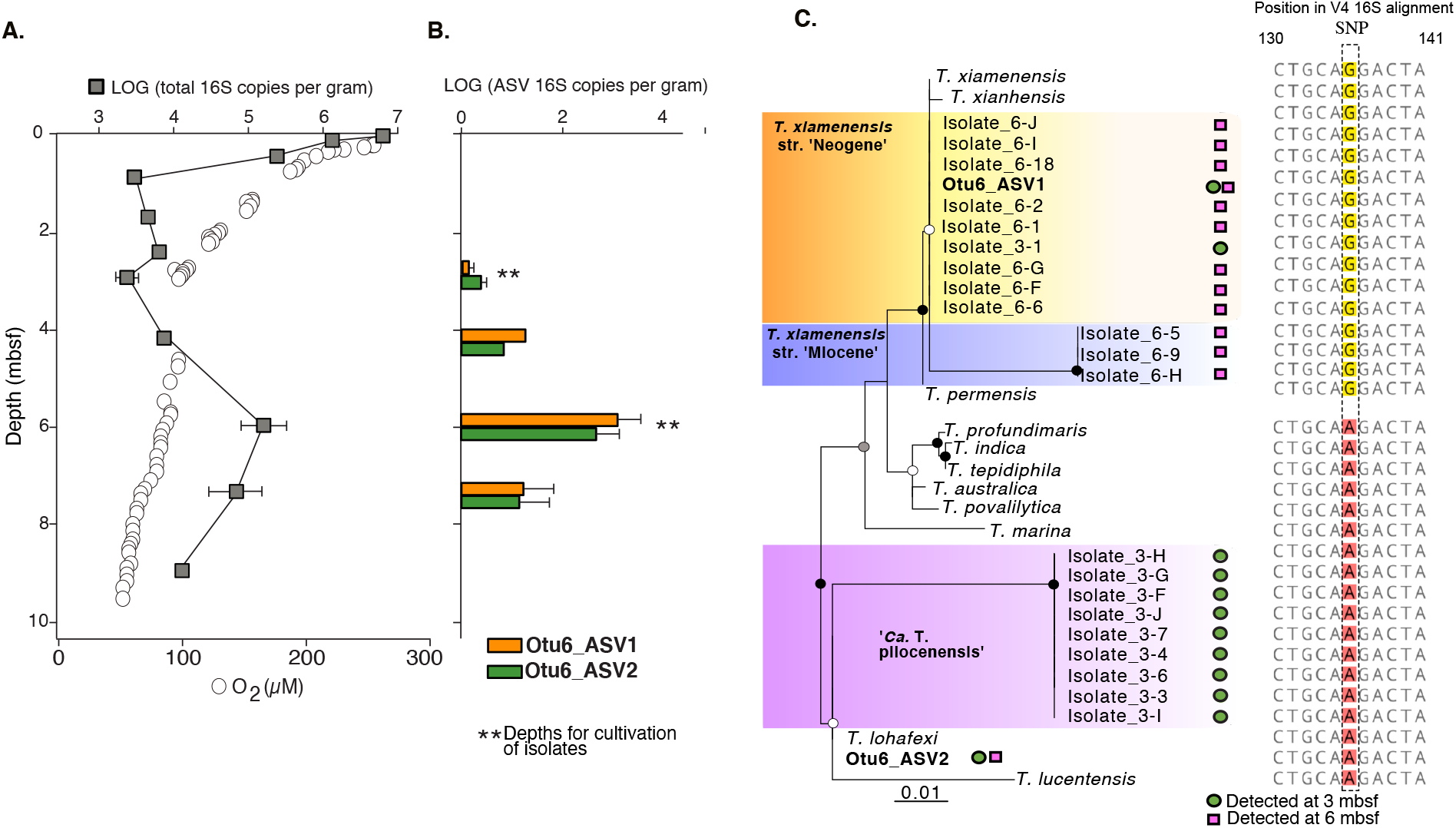
Isolated subsurface *Thalassospira* are most abundant at 3-7 mbsf and distinct from related type strains isolated from overlying water and sediments. (A) Vertical profile of total 16S rRNA gene concentrations determined via qPCR (squares), and oxygen concentrations (circles). 16S rRNA gene concentration points are the average abundances of three technical qPCR replicates with ranges shown with error bars. (B) qPCR-normalized average concentrations of the *Thalassospira* affiliated ‘Otu6’ ASVs. Error bars are ranges from three technical qPCR replicates. Asterisks mark the depths for the long term ^18^O-water incubation experiments, enrichments, and cultivation. (C) Maximum likelihood (PhyML) phylogenetic analysis of the *Thalassospira* 16S rRNA gene ASVs (V4 hypervariable region), together with subseafloor and type strain *Thalassospira* 16S rRNA gene V4 regions. The presence of the SNP is displayed. Black, grey, and white dots at the nodes represent >90%, >70%, >50% bootstrap support, respectively.

We isolated colony-forming bacteria on petri dishes following an 18-month incubation of sediment and sterile ^18^O-labeled artificial seawater (Fig S1) from 3 and 6 meters below the seafloor (mbsf)(see Methods). Because the mean sedimentation rate is 1 m million yr^−1^, the age of the sediment horizons from which these bacteria were enriched and isolated are estimated to be 3 and 6 million years old, respectively. The full-length 16S rRNA gene sequences from the isolates had closest similarity (90-99% sequence identity) to the Alphaproteobacteria *Thalassospira xiamenensis and Thalassospira lohafexi* previously isolated from marine surface sediment (*16, 17*) and oligotrophic seawater (*18*) (these previously isolated microbes and their related cultured relatives are referred to as ‘type strains’ herein).

Several lines of evidence indicate the *Thalassospira* isolated from the 3- and 6-mbsf sediment enrichments are endemic to the subseafloor clay and are not a contaminant from the water column or other sources. First, the V4 hypervariable region of the 16S rRNA gene sequences from the sediment slurry enriched *Thalassospira* cultures share >99% sequence identity with an operational taxonomic unit (OTU) previously identified from the *in-situ* community determined from the frozen samples (‘OTU_6’, Figure 1B,C). This OTU_6 became ^18^O-labeled during the 18-month enrichment incubation (atomic ^18^O enrichment of OTU_6 DNA: 59%) in the presence of sterile ^18^O-labeled seawater (Fig. S1), a proxy for growing microbes (*19*), and had an estimated doubling time in the incubation of 36 ± 1.5 (mean ± SD) days. OTU_6 consists of two amplicon sequence variants (ASVs) that cluster with the *Thalassospira* subseafloor isolates (Otu6_ASV1, Otu6_ASV2), respectively, and are distinguished by a single nucleotide polymorphism (SNP) that is conserved between the ASVs and the subseafloor isolates (Figure 1C). The *insitu* concentrations of both *Thalassospira* ASVs have highest abundance (ca. 1,000 16S rRNA gene copies g^−1^ sediment) below the seafloor between 4 - 6 mbsf, and both *Thalassospira* ASVs were detected in the 3- and 6-mbsf sediment (Figure 1B). This shows that the *Thalassospira* strains isolated from the 3- and 6-mbsf sediment enrichments are derived from the same distinct 16S rRNA gene ASVs present within the *in-situ* communities. The long-term physical isolation of these isolates in the subseafloor (see ‘sediment physical properties’ in SI), subsisting under uninterrupted energy limitation within the ancient sediment, provides an opportunity to investigate how the relative effects of recombination, nucleotide substitution, and gene decay have shaped the genomes of the cultivated subsurface *Thalassospira* strains since their burial in the deep-sea clay millions of years ago.

### Genome statistics

We sequenced the genomes of ten *Thalassospira* isolates each from the 3- and 6-mbsf sediment enrichments using a hybrid assembly approach consisting of long-read Nanopore sequencing, corrected and polished via short-read Illumina technologies at >100x coverage. The mean genome completeness of these hybrid assemblies is estimated to be 99.7% ± 0.3%; (mean ± SD), with most being 100% complete and representing the complete chromosome (Table S1). The mean length of the new *Thalassospira* genomes is 4.71 ± 0.08 Mbp (mean ± SD), with 4,567 ± 107 (mean ± SD) protein coding genes, and they are assembled to an average of 12 ± 2 (mean ± SD) contigs (Fig. S2 and Table S1). The genome size and the number of protein-encoding genes are similar to those observed within the existing *Thalassospira* type strains isolated from the surface world (Fig. S2).

### Core genome phylogenomic analysis

The core genome phylogeny of existing *Thalassospira* type species, and the newly isolated subseafloor *Thalassospira* strains, consists of 1,809 orthologous genes and reveals three clades of subseafloor *Thalassospira*. One clade shares 96-97% genome-wide average nucleotide identity (ANI), and 99.9% 16S rRNA gene sequence identity, with *T. xiamenensis* and *T. permensis* (Fig. S3). We named isolates in this clade *T. xiamenensis* strain ‘Neogene’, after the Neogene eon (2.8 – 23 mya) which covers both estimated ages of sediment (3 mya and 6 mya) from which the strains in this clade were isolated. The subseafloor genomes in this clade correspond to the 16S rRNA gene ASV1 detected in the *in situ* frozen sediment core samples (Fig 1B, C). A second clade contains three isolates from 6 mbsf sediment shared 97% ANI with *T. xiamenensis* and *T. permensis*. Since all isolates in this clade were recovered from ca. 6 mya sediment deposited during the Miocene eon (5.33 – 23 mya), we report them as *T. xiamenensis* strain ‘Miocene’ (Fig. S3). However, despite sharing 97% ANI in the core genome with *T. xiamenensis* and *T. permensis* (Fig S3), *T. xiamenensis* strain ‘Miocene’ only shared 90% 16S rRNA gene sequence identity with these closest related type strains. The subseafloor genomes in this clade also correspond to ASV1 detected in the *in situ* frozen sediment core samples (Fig 1B, C). A third clade contains subseafloor *Thalassospira* cultures isolated only from 3 mbsf sediment and shares 95-96% ANI with *T. lucentensis* and *T. lohafexi* (Fig. S3). The subseafloor genomes in this clade correspond to the 16S rRNA gene ASV2 detected in the *in situ* frozen sediment core samples (Fig 1B,C). Based on the genetic distinctness of these isolates we consider them to be a new candidate species, according to recently provided criteria based on genome-wide ANI (*20*). Because all isolates of this third clade were recovered from 3 mya sediment, we propose the candidate name *‘Candidatus* Thalassospira pliocenensis’, named after the Pliocene age (2.58 – 5.33 mya) of the deep-sea clay from which they were isolated. Pangenome analysis revealed that flexible genome content is conserved within each of the three subseafloor clades, providing further evidence that each clade represents a genetically distinct population (Fig S4). Laboratory growth rates of the subseafloor *Thalassospira* ranged from 0.064 to 0.31 h^−1^ (Fig. S5).

### Roles of mutation and recombination

The ratio of nucleotide substitutions originating from mutations versus homologous recombination (r/m) can be used to measure the relative effect of homologous recombination on the genetic diversification of populations (*21*). Due to the physical isolation of individual bacterial cells, reduced cell concentrations, and the reduced availability of extracellular DNA for recombination in subseafloor sediments (*22*) we hypothesized that r/m ratios would be lower in the subseafloor *Thalassospira* populations than in the type strains. To test this, we used an established method (*23*) to calculate the relative rate of recombination to mutation (*R/θ*), the mean length of recombined DNA (*δ*), and the mean divergence of imported DNA (*v*) for branch tips (existent genomes) and internal nodes (ancestral states) in the *Thalassospira* core genome phylogeny, which allows for a calculation of r/m (r/m = (*R/θ*) * *δ* * *v*). This analysis showed that in the *Thalassospira* core genome, the r/m is approximately ten times lower in the subseafloor core genome (r/m = 0.078) than in the type strains (r/m = 0.71) (Table 1), indicating that homologous recombination plays a much lesser role in the diversification of the subseafloor strains. The r/m values of the subseafloor *Thalassospira* are furthermore anomalously low compared to free-living bacteria isolated from the surface world, which have r/m values that range from 0.1-64 (*21*). Concomitant with the ten-fold lower r/m values compared to the type strains (Table 1), the subseafloor *Thalassospira* core genomes exhibit far fewer numbers of inferred imported DNA from recombination events compared to the *Thalassospira* type strains and the ancestral states of the last common ancestors shared with the type strains (Fig. 2).

**Table 1.** The contributions of recombination and mutation to nucleotide diversity in subseafloor populations. The results from ClonalFrameML (*23*) analysis used to calculate the relative contributions of recombination and mutation in the core genome (r/m). *R/θ*: the relative rate of recombination compared to mutation, *δ*: the average length of recombined (imported) DNA, *v*: mean divergence of imported DNA. Also displayed are the average number of pseudogenes and dN/dS ratios (+/- standard deviation). ** two sided T-test: P = 0.005. *****two sided T-test: P = 0.000001.

| Group | # strains | $R/\theta$ | $\delta$ | $\nu$ | $r/m$ | # pseudogenes | dN/dS |
| --- | --- | --- | --- | --- | --- | --- | --- |
| All subseafloor and type strains | 34 | 0.053 | 244 | 0.055 | 0.71 | 37 (+/- 8) | 0.025 (+/- 0.011) |
| Type strains | 13 | 0.04 | 333 | 0.053 | 0.71 | 21 (+/- 3) | 0.022 (+/- 0.01) |
| Subseafloor | 21 | 0.006 | 500 | 0.026 | 0.078 | 48 (+/- 4)**** | 0.038 (+/- 0.007)** |

**Figure 2:**
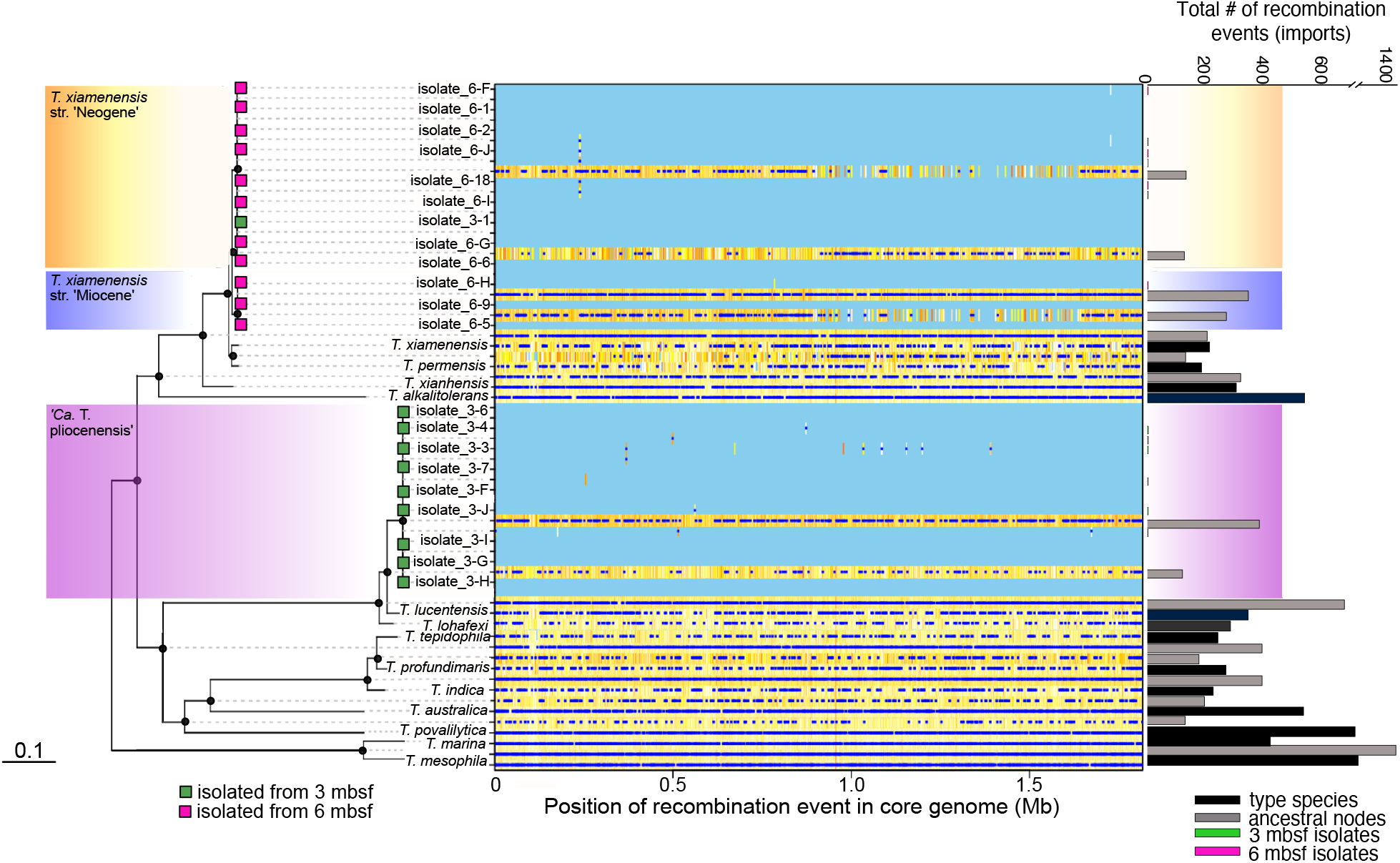
Recombination in the conserved core genome is limited in subseafloor *Thalassospira* populations. The maximum likelihood (PhyML) phylogenetic tree is based on a concatenated alignment of 1,809 genes conserved across all *Thalassospira* genomes (‘core genes’). Black circles on nodes represent bootstrap values >95%. The position of recombination events in the core genome are represented by dark blue dots. Positions of low nucleotide diversity and no recombination events in the core genome are shown in light blue. Nucleotide diversity at specific sites in the core genome are illustrated with a color gradient (white: less diversity, orange: more diversity). Histograms on the right display the total number of recombination events (imports) in each genome sequence, and ancestral state reconstructions (internal nodes), as detected by ClonalFrameML (*23*).

The ancestral recombination signal is not preserved in the genomes of the subseafloor isolates (Fig. 2), which is consistent with a shift away from a recombination-influenced population toward a more clonal population structure since the subseafloor *Thalassospira* diverged away from the last common ancestor shared with the type strains. We looked for evidence of evolution by investigating pairwise substitution numbers in the subseafloor *Thalassospira* genomes. We identified tens to thousands of nucleotide differences (single nucleotide polymorphisms [SNPs]) within each subseafloor *Thalassospira* clade (Fig S6). The SNPs in subseafloor *Thalassospira* strains were present in a clade-specific manner (Fig S7). But, while the precise SNPs are clade-specific, the SNPs occurred in genes coding for related functions across different clades. These included genes with predicted annotations involved in flagellar motility (FlhB, FliO), transcription (TetR and Fis family transcriptional regulators), cell wall biogenesis (peptidase S41, peptidoglycan DD-metalloendoptidase M23), and transport and metabolism of amino acids and carbohydrates (Fig. S7).

We considered that the observed nucleotide substitutions occurred within the subseafloor populations during the culture enrichment and cultivation process. However, the measured generation times of the *Thalassospira* (OTU6) in the incubation measured with qPCR (36 ± 1.5 days; mean ± SD) indicate an estimated maximum of 15 doublings over the enrichment (see SI). At common bacterial mutation rates of 10^−9^ to 10^−10^ mutations bp^−1^ generation^−1^ (*24*), more than 4,000 generations would be required to obtain the observed nucleotide diversity present in the subsurface *Thalassospira* genomes (Fig. S8). Thus, newly accumulated mutations in culture are insufficient to explain the observed interpopulation nucleotide diversity between the subseafloor genomes (Fig. S8). The inter-population nucleotide diversity of the subseafloor strains therefore arose during their long-term subsistence in the ancient sediments and is not the result of evolution during the laboratory incubation.

### Substitutions and pseudogenes are fixed in subseafloor populations

Compared to the type strains, the subseafloor *Thalassospira* genomes exhibit higher numbers of pseudogenes (non-functional parts of the genome that resemble functional genes), and non-synonymous substitutions (substitutions that alter the amnio acid sequence of a protein). We identified 47.9 ± 8.57 (mean ± SD) pseudogenes in the genomes of subseafloor *Thalassospira* isolates, which is significantly higher than the number of pseudogenes identified in the type strains (22.1 ± 5.52 pseudogenes [mean ± SD]; Table 1, Figure 3, and Fig. S2) (two-sided T-test: P=1.5E-10). Similarly, we observed a modest but significant elevation of genome-wide nonsynonymous to synonymous substitution rates (dN/dS) in the genomes of the subseafloor *Thalassospira* strains (0.035 ± 0.006; mean ± SD), relative to the type strains (0.022 ±0.012; mean ± SD) (two-sided t-test P=0.0002; Table 1, Figure 3 and Fig. S2). Similar to the SNPs (Fig. S7), the composition of pseudogenes occurred in a clade-specific manner (ANOSIM: P = 0.001) (Fig S9). Compared to the type strains, the predicted annotations of the pseudogenes in the subseafloor *Thalassospira* genomes are skewed toward those involved in transcription, energy conservation, amino acid and carbohydrate metabolism, and flagellar motility (Fig S9). Moreover, subseafloor genomes have significantly higher numbers of pseudogenes involved in motility (flagellar biosynthesis genes: FliN, FliK, FlhO) compared to the type strains (two-sided t-test: *P*=0.003) (Fig S10).

**Figure 3:**
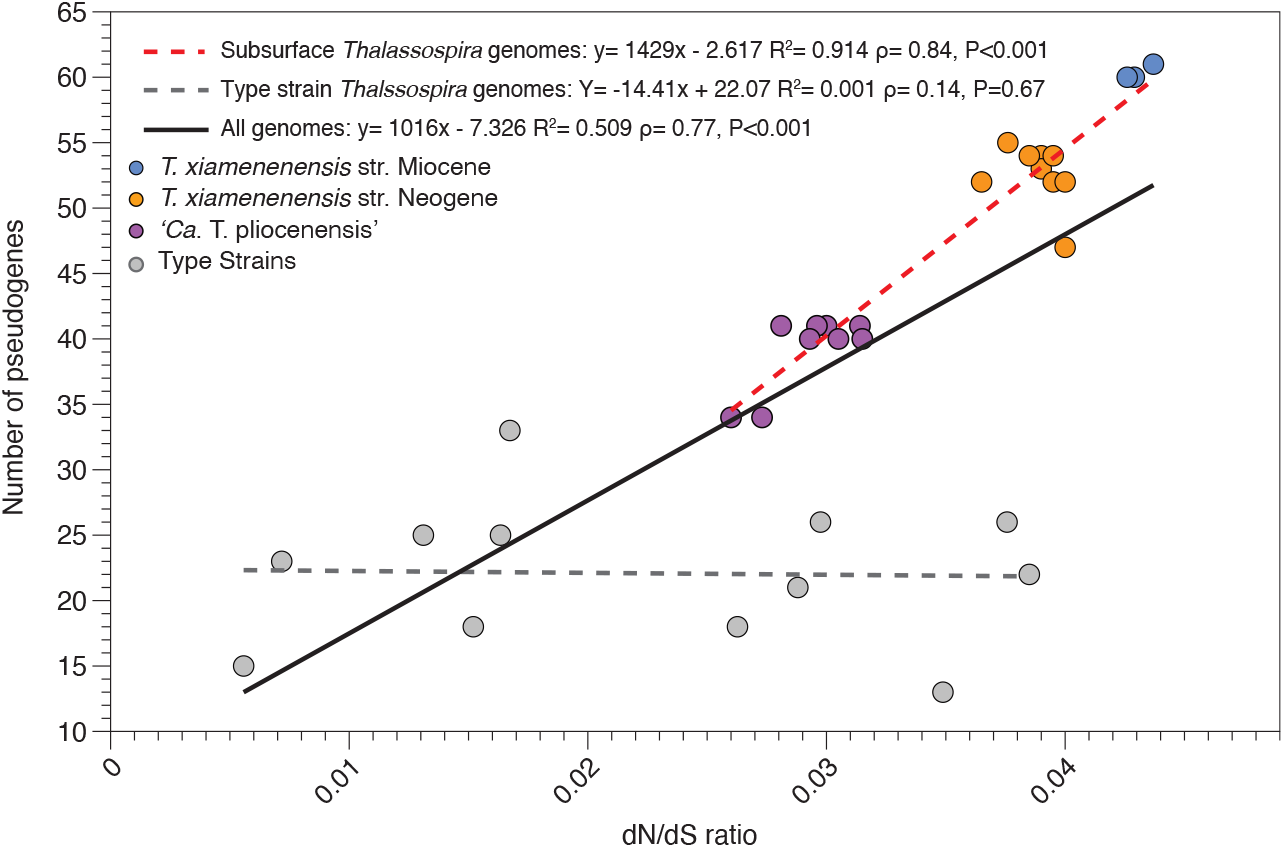
The number of pseudogenes and dN/dS ratios are elevated and correlated in subseafloor *Thalassospira* populations. Subseafloor genomes accumulate more pseudogenes as a function of increasing dN/dS ratios compared to the type strains. Linear regressions for type strains, subseafloor strains, and all strains are displayed.

The elevated and correlated dN/dS ratios and proportion of pseudogenes for these isolates (Fig. 3) is a hallmark of Muller’s ratchet (*24–28*) and is a further indication that the three clades of subsurface *Thalassospira* are clonal and unable to eliminate deleterious mutations that otherwise would be purged from natural populations that are able to freely recombine. Burial in the sediment millions of years ago resulted in physical isolation of subsurface *Thalassospira* cells and was the impetus for a transition from freely recombining populations to recombination-limited clonal populations that rarely encounter genetically diverse recombination partners that introduce genetic diversity into the population. Simultaneously, the effective population sizes of subsurface *Thalassospira* were restricted by the very low flux of bioavailable energy in subseafloor sediment. The reduced energy flux limited the environmental carrying capacity (*1*), and thus the *Thalassospira* population size, in this ancient subseafloor clay. The lack of available energy also likely significantly reduced the potential for cellular motility (*5, 22*). The combined effects of physical isolation, low cell concentrations, and little or no motility, resulted in the further reduction of recombination events due to infrequent cell-cell contact. The reduction of homologous recombination resulted in mutation becoming the dominant driver of evolution in these clonal subseafloor *Thalassospira* populations (Table 1), leading to relaxed purifying selection in the absence of recombination. Although we see elevated dN/dS ratios and an accumulation of pseudogenes across the genomes of subsurface *Thalassospira* (Fig. 3), some functions appear to be more prone to gene decay than others. For example, genes predicted to be involved in flagellar motility were present in both the SNP (*fliO, flhB, flgH*; Fig. S7) and pseudogene (*fliN, fliK, flhO*; Fig. S9, S10) analysis, suggesting flagellar motility may become a non-essential trait (*5, 22*) subject to purifying relaxed selection in abyssal clay with (which is characterized by very low permeability and extremely small pore diameter, despite its high porosity [*15*]). Finally, the accumulation of mutations via Muller’s ratchet (*24*) may explain the reduced growth rates (29) of the subsurface *Thalassospira*, relative to the type strains (Fig. S5).

### Outlook

Our findings demonstrate that subseafloor *Thalassospira* genomes analyzed here have evolved akin to endosymbiotic bacteria, whereby clonal populations also lack homologous recombination and are thus subject to genetic drift whereby deleterious mutations become fixed and Muller’s Ratchet (*24*) eventually leads to the extinction of endosymbiotic bacterial lineages (*25*). Our genomes show similar signs of evolution, but dN/dS ratios observed in the subseafloor *Thalassospira* genomes are lower than those seen in endosymbiotic bacteria (*28*) and genome reduction was absent. This could be explained by the subseafloor genomes experiencing a single bottleneck in the form of a burial event followed by a stable but low population size, in contrast to the repeated population bottlenecks experienced by endosymbionts at each insect generation.

Subseafloor cell concentrations are low and decrease substantially with increasing depth (*1*). Because genetic drift has a stronger effect on populations with small population sizes (*25, 28*), physically isolated microbes experiencing clonal growth and reduced homologous recombination in the deep biosphere may be particularly prone to genetic drift-mediated evolution. Because the subseafloor biosphere contains a large fraction of all bacterial cells on Earth (*1*), our findings suggest drift-like evolutionary processes in the absence of homologous recombination may be much more widely distributed in nature than previously thought. Future assessments of homologous recombination and drift in single cell genomes from uncultured lineages of bacteria and archaea that comprise most subsurface energy limited communities (*30*) could be used to assess how widespread this evolutionary mechanism is within the subsurface biosphere.

## Acknowledgements

This work was supported primarily by the Deutsche Forschungsgemeinschaft (DFG) project OR 417/1-1 granted to W.D.O. Publication of the manuscript was supported by the LMU Mentoring Program. The expedition was funded by the US National Science Foundation through grant NSF-OCE-1433150 to S.D. This is Center for Dark Energy Biosphere Investigations (C-DEBI) publication number XXX. A portion of this work was performed as part of the LMU Masters Program “Geobiology and Paleobiology” (MGAP). P.C. was supported by the University of Arizona’s Technology and Research Initiative Fund (the Water, Environmental, and Energy Solutions initiative). We thank William F. Martin for comments on the manuscript.

## Author contributions

W.D.O., P.C., J.S., and G.W. conceived the work and experimental approach. W.D.O., T.M., O.K.C., S.V., P.C., S.H., and A.V. contributed to the laboratory and bioinformatics analyses and experimental work. S.D. provided the samples from the KN223 R/V Knorr oceanographic expedition KN223. All authors discussed and wrote the manuscript and commented on the paper.

## Supplemental information

Supplemental Table 1, Supplemental Figures S1–S10, References 31-49.

## Data and materials availability

Data are publicly available through NCBI BioProject PRJNA473406. The raw genomic sequences reads and genome assemblies are available in SRA BioSample accessions SAMN17168194, SAMN17168195, and SAMN17168196. The 16S data from the frozen core are available in SRA BioSample accessions SAMN10929403 to SAMN10929517. Figures and output files from the pangenomic analysis in Anvio are available online through FigShare (https://doi.org/10.6084/m9.figshare.13372619). Additional data related to this paper may be requested from the authors.

## Supplemental Information

**Table S1.**
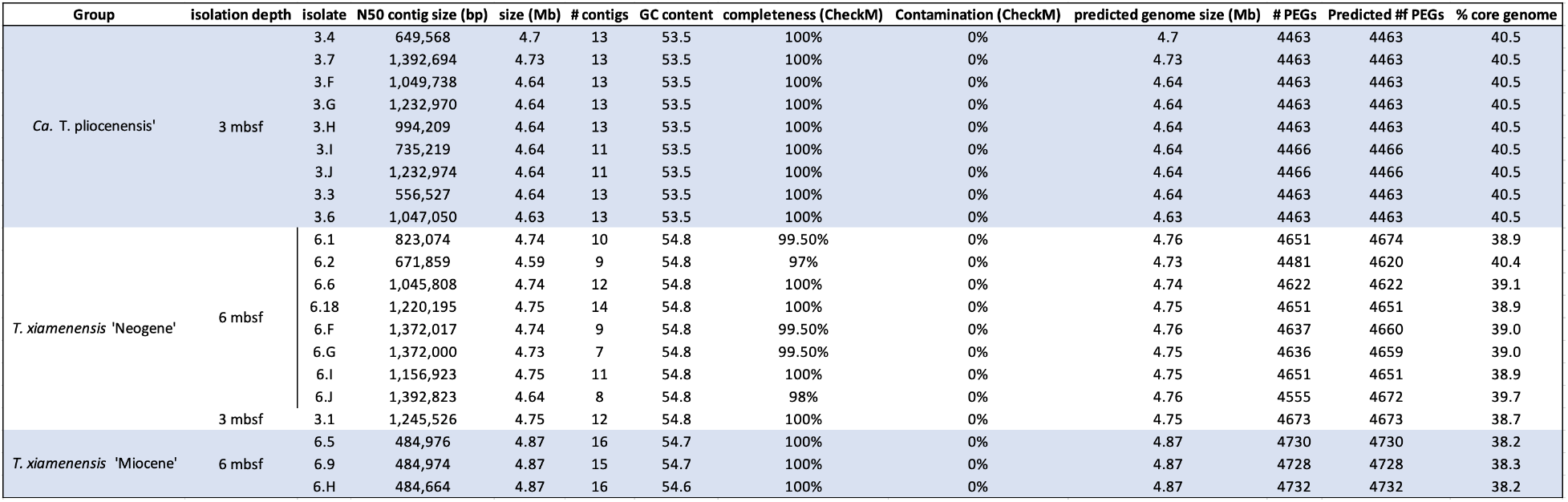
Genome summary statistics.

**Figure S1.**
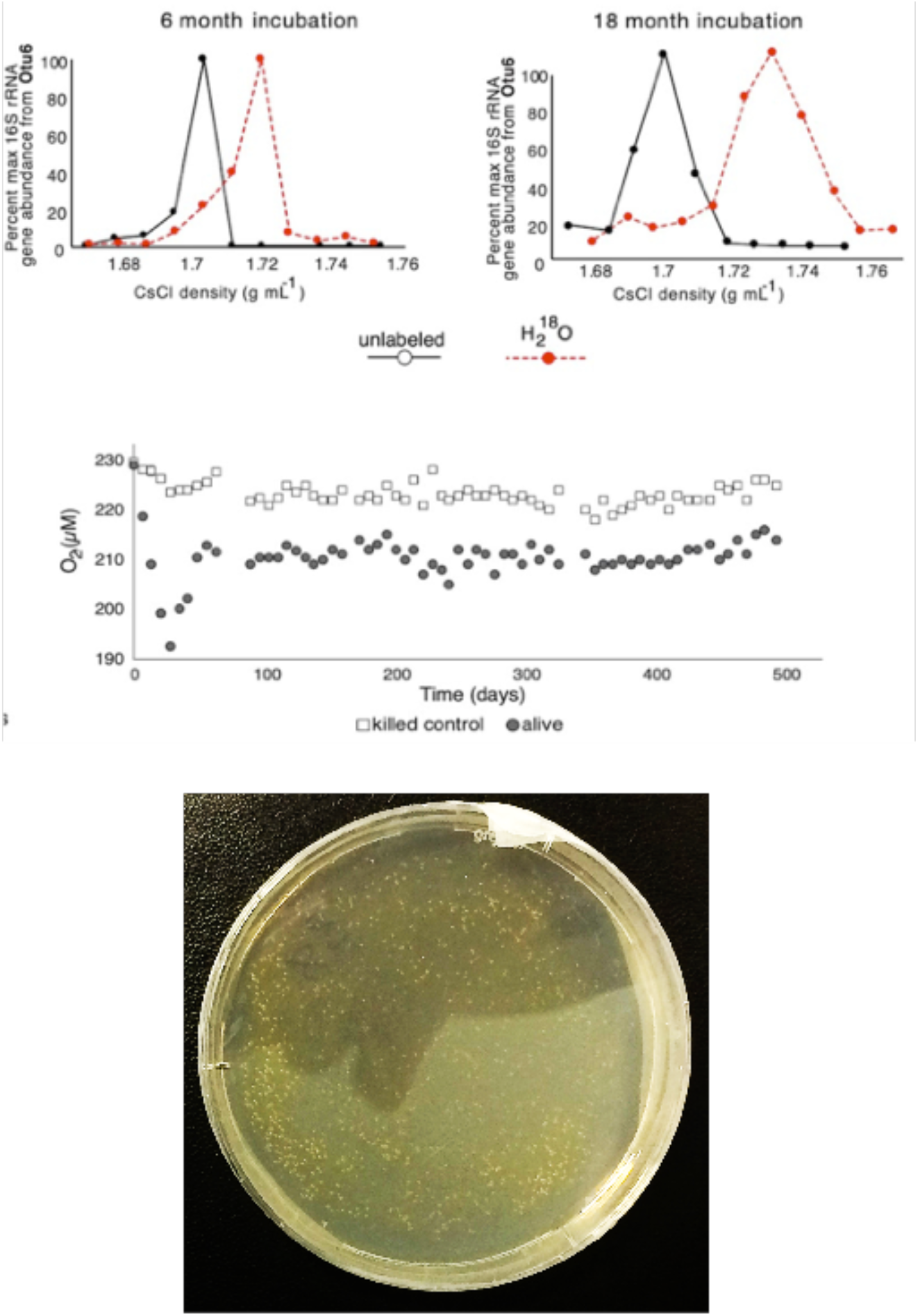
**Top panel:** ^18^O-labeling of 16S rRNA genes from the *Thalassospira* OTU6 (see Figure 1), after 7 and 18 months of incubation with ^18^O-labeled water from the 3 mbsf sediment (data originally published in Vuillemin et al., 2019). **Middle panel:** Oxygen consumption over time in the 18 month slurry from the 3 mbsf incubation (filled circles), and slurries containing labeled water and autoclaved sediment (killed control). **Bottom photo**: cultivation of colony forming bacteria on solid media after the 18 month incubation of sediment slurries in sterile ^18^O-labeled artificial sea water. No bacterial colonies formed on petri dishes that were inoculated with the killed control slurries.

**Figure S2:**
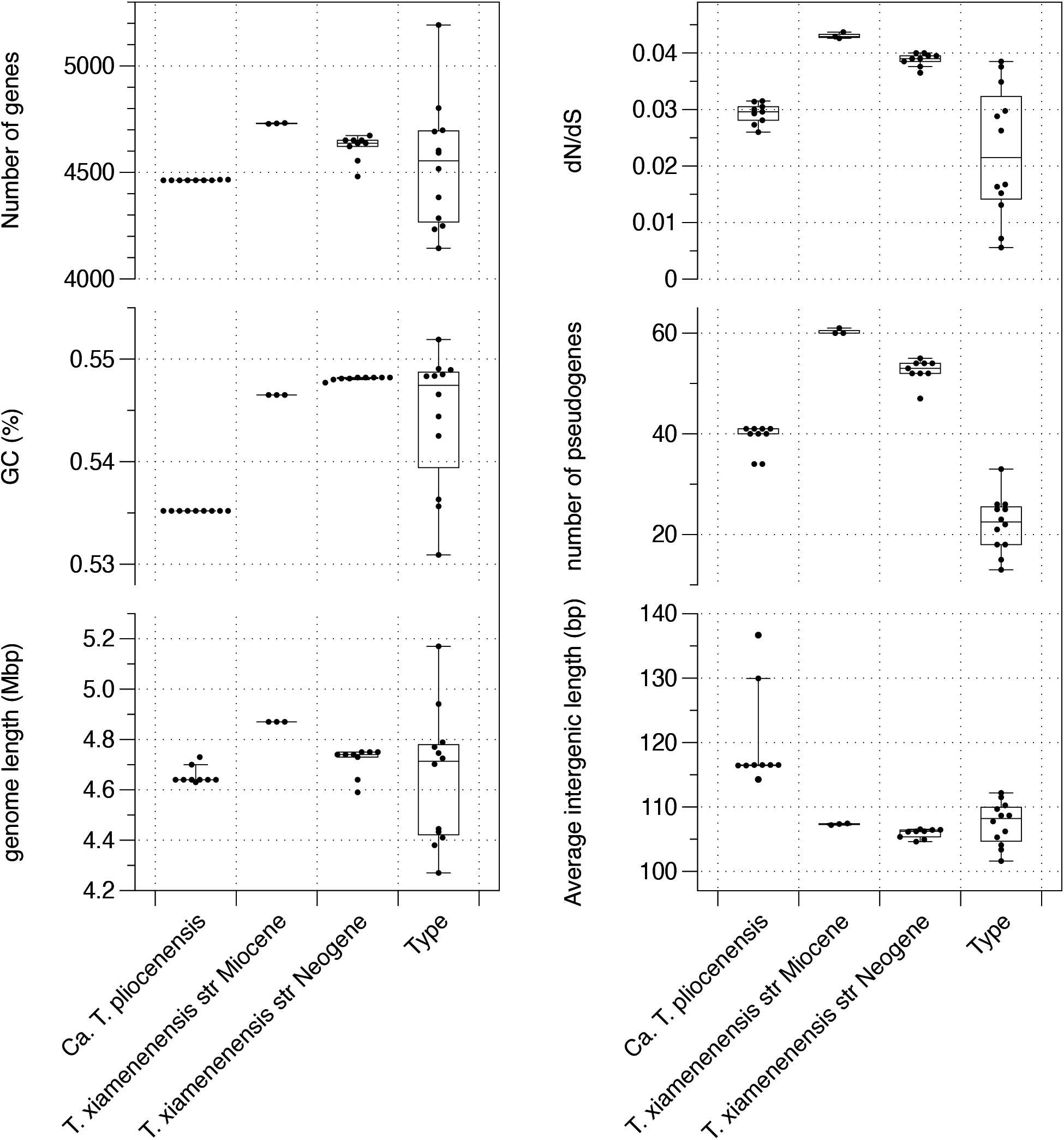
Summary of genome properties for *Thalassospira* strains used in this study. Points are derived from the analysis of existing genome sequences (for “Type” strains), and new high-quality draft genomes sequenced as part of this study. Box plots illustrate interquartile range ± 1.5 × interquartile range. The horizontal line in each box plot is the median.

**Figure S3.**
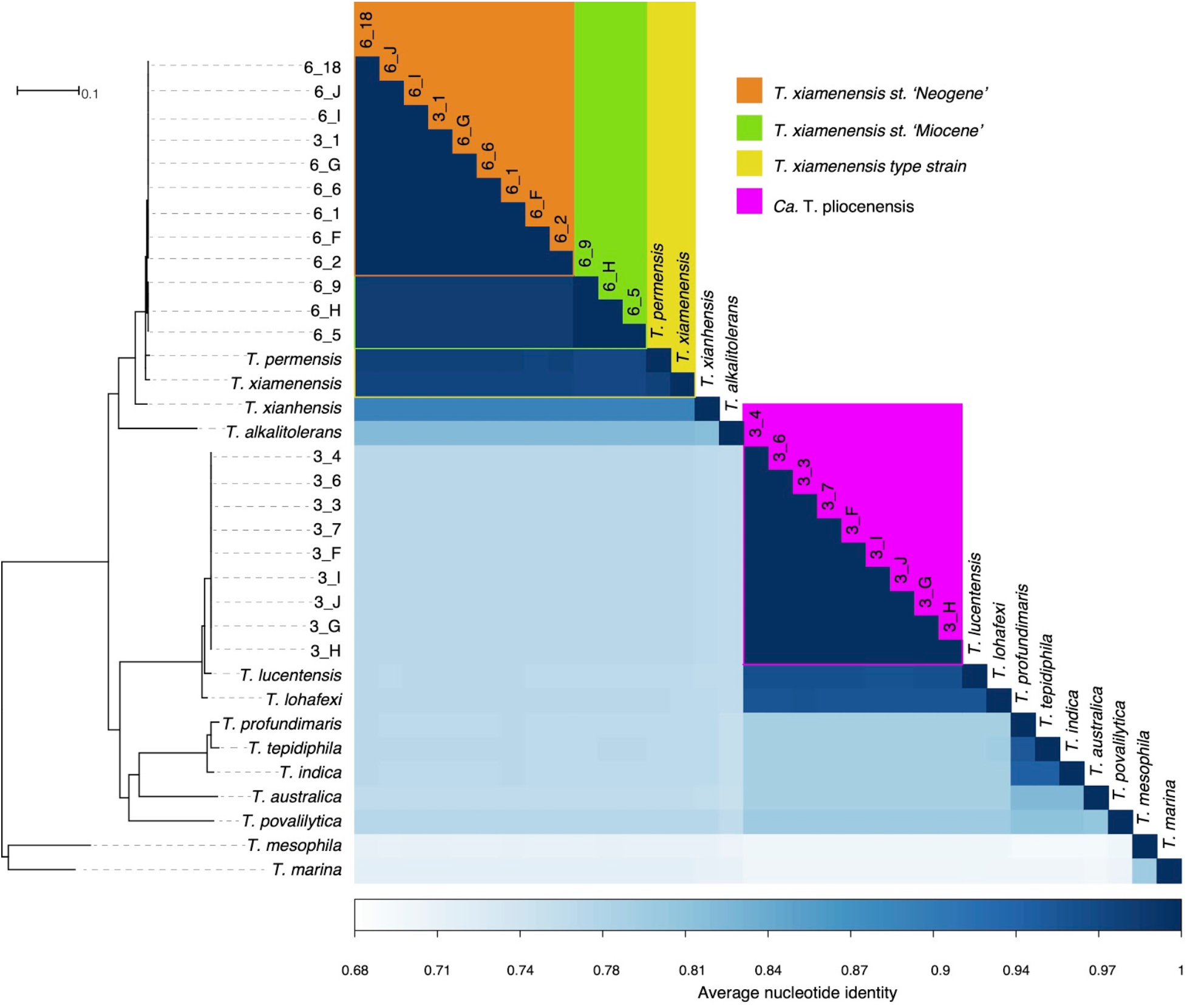
Average nucleotide identity (ANI) and the core genome phylogeny. The tree is based on maximum likelihood and a concatenated alignment of 1,809 core genes

**Figure S4:**
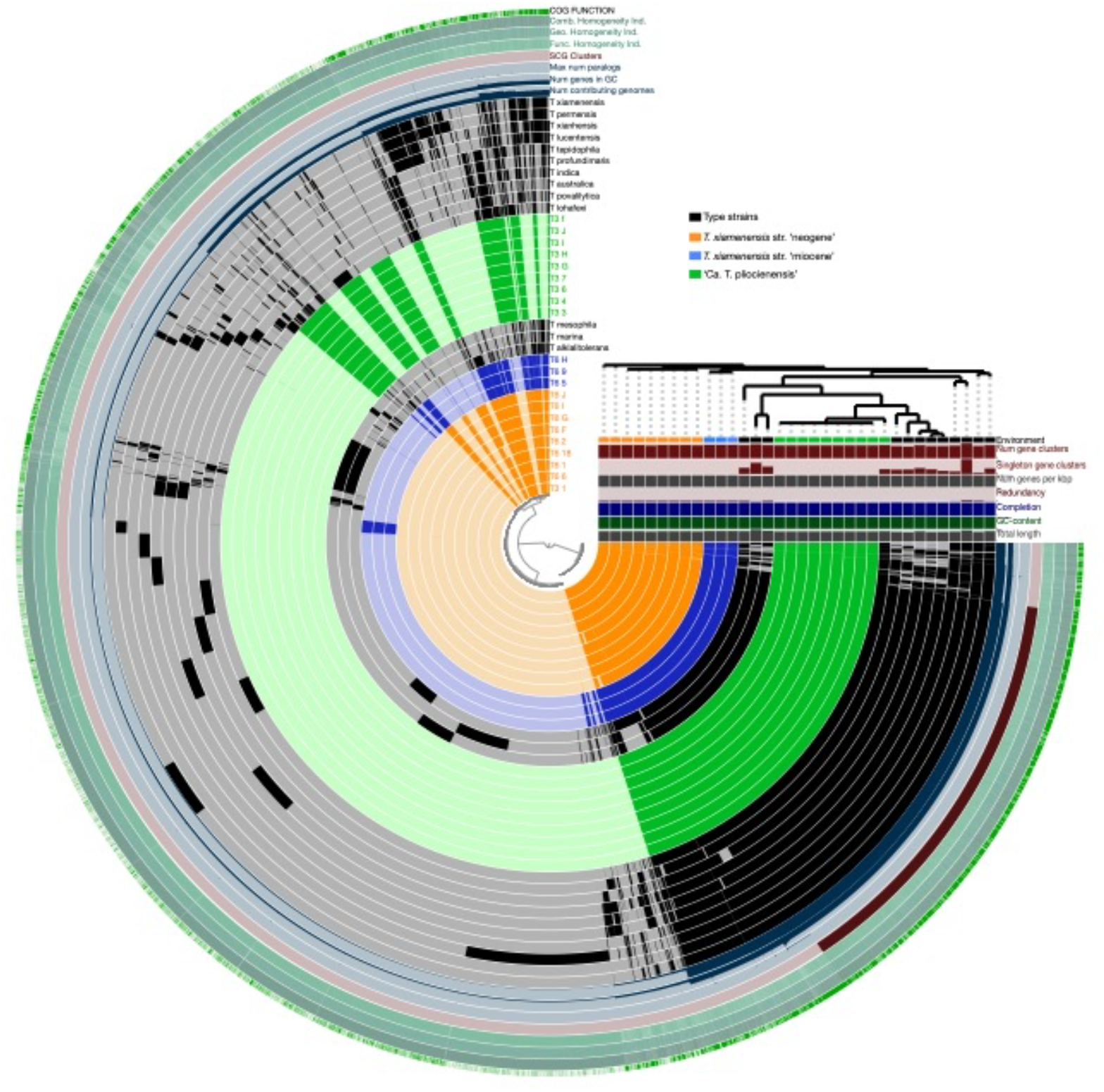
Pangenome analysis of all *Thalassospira* genomes included in the study. The internal dendrogram is a UPGMA based on the presence/absence of shared gene orthologs. Black bars in the first (inner) 34 circles show the occurrence of gene clusters in the genome of *Thalassospira* species. Grey areas and light colors in the circles represent gene clusters that were detected in the corresponding genome. The next eight bars show statistics for the pangenome analysis of each individual gene cluster (inner circle to outer circle) # contributing genomes: # of genomes that has a hit in a gene cluster, (GC), max # paralogs, single copy gene clusters (SCG), Functional Homogeneity index, Geometric homogeneity index, combined homogeneity index, presence of a COG functional assignment. The categories on the right side (below dendrogram) show the totals per genome for # of gene clusters found in each genome, Num genes per kbp: Number of genes per kilobase pairs of genome. Redundancy: Multiple occurrence of single copy genes in a genome, Completeness: Calculated from the occurrences of single copy gene set in a genome

**Figure S5.**
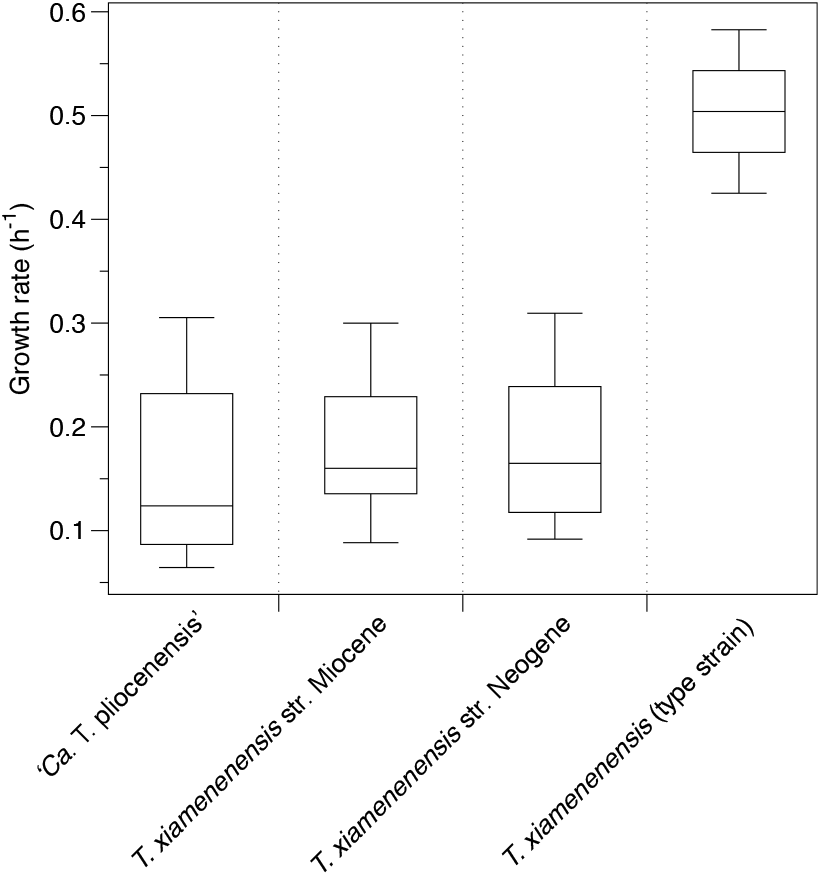
Laboratory growth rates of subsurface *Thalassospira* isolates in comparison to type strain *T. xiamenenensis*. Box plots illustrate interquartile range ± 1.5 × interquartile range. The horizontal line in each box plot is the median. For subsurface strains, growth rates were calculated for each strain across three independent experiments. For the *T. xiamenenesis* type strain, growth rates were calculated from triplicate flasks of one experiment.

**Figure S6.**
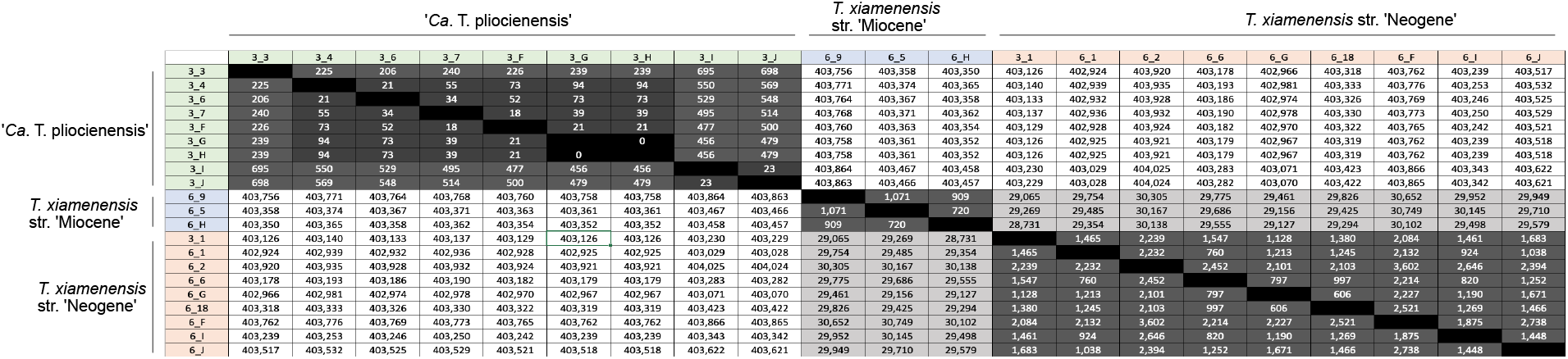
The number of SNPs between pairs of subseafloor *Thalassospira* genomes.

**Figure S7.**
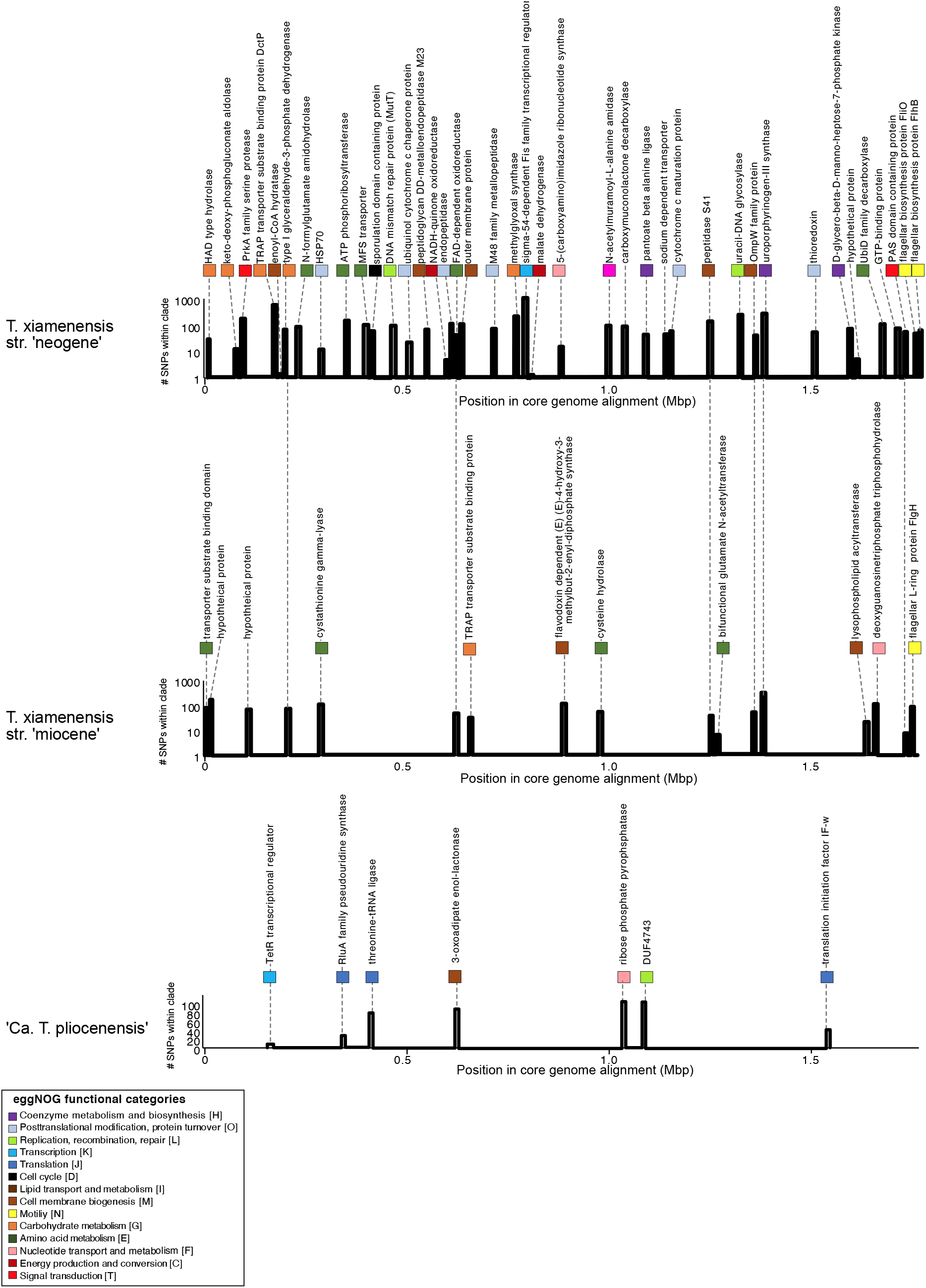
The number of interpopulation SNPs at different positions in the core genome alignment, for each of the three subseafloor populations. The gene annotations to the corresponding regions are shown.

**Figure S8:**
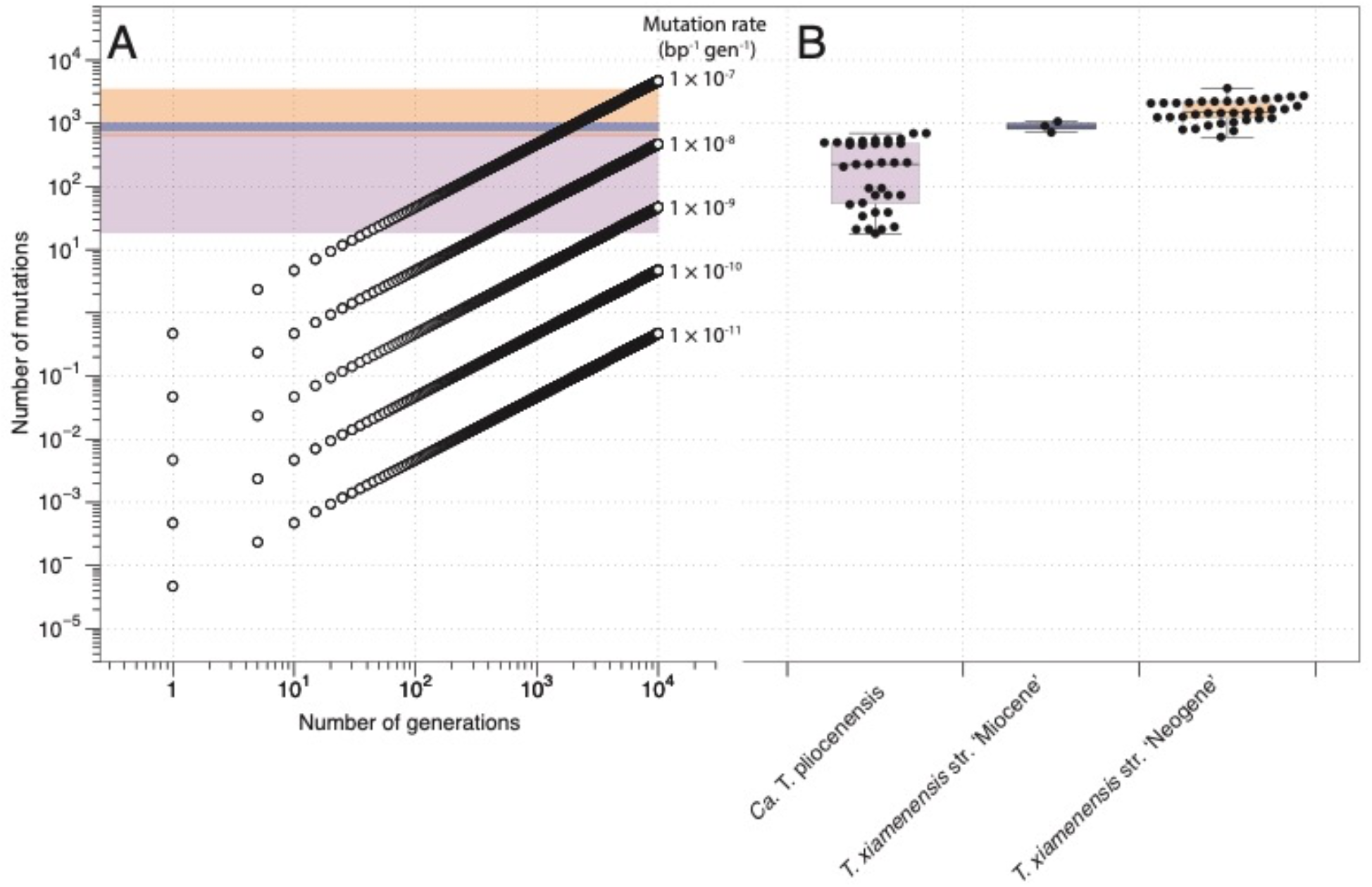
Interpopulation nucleotide diversity did not arise during laboratory cultivation. (A) Model of the number of mutations expected per genome (4.7 Mbp) as a function of total number of generations at five different mutation rates. Shading depicts the range of observed nucleotide diversity in subseafloor *Thalassospira* genomes (number of pairwise SNPs) as described in panel B. (B) The distribution of interpopulation nucleotide diversity. Points are the number of pairwise SNPs detected in subseafloor *Thalassospira* genomes. Box plots illustrate interquartile range ± 1.5 × interquartile range. The horizontal line in each box plot is the median.

**Figure S9:**
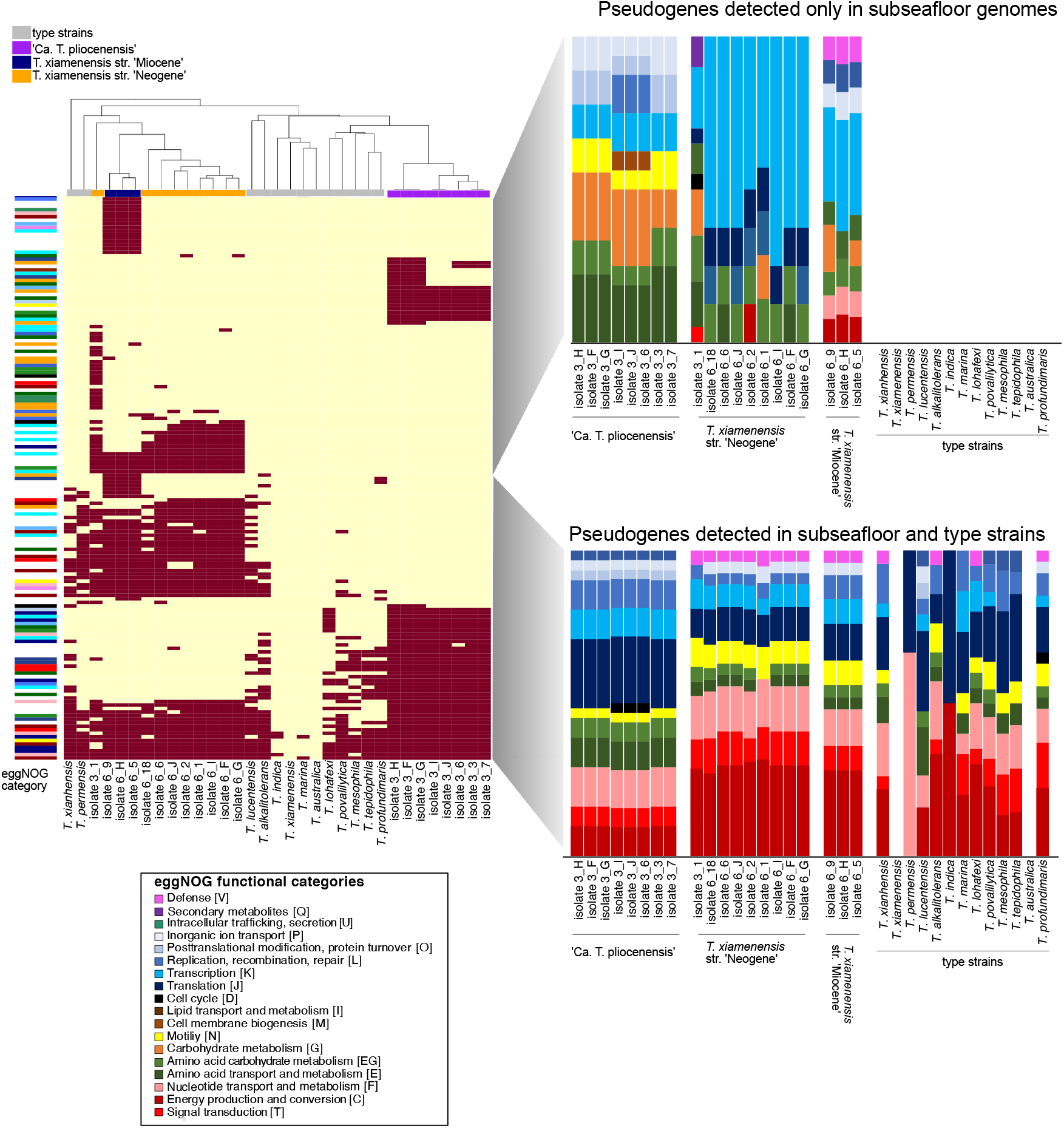
The heatmap shows the presence/absence of pseudogenes in the subseafloor genomes and the presence of these pseudogenes in type; strains. Functional annotation (against eggNOG) of pseudogenes found in the subseafloor genomes only are compared to functional annotations of pseudogenes found in both the subseafloor and type strains.

**Figure S10:**
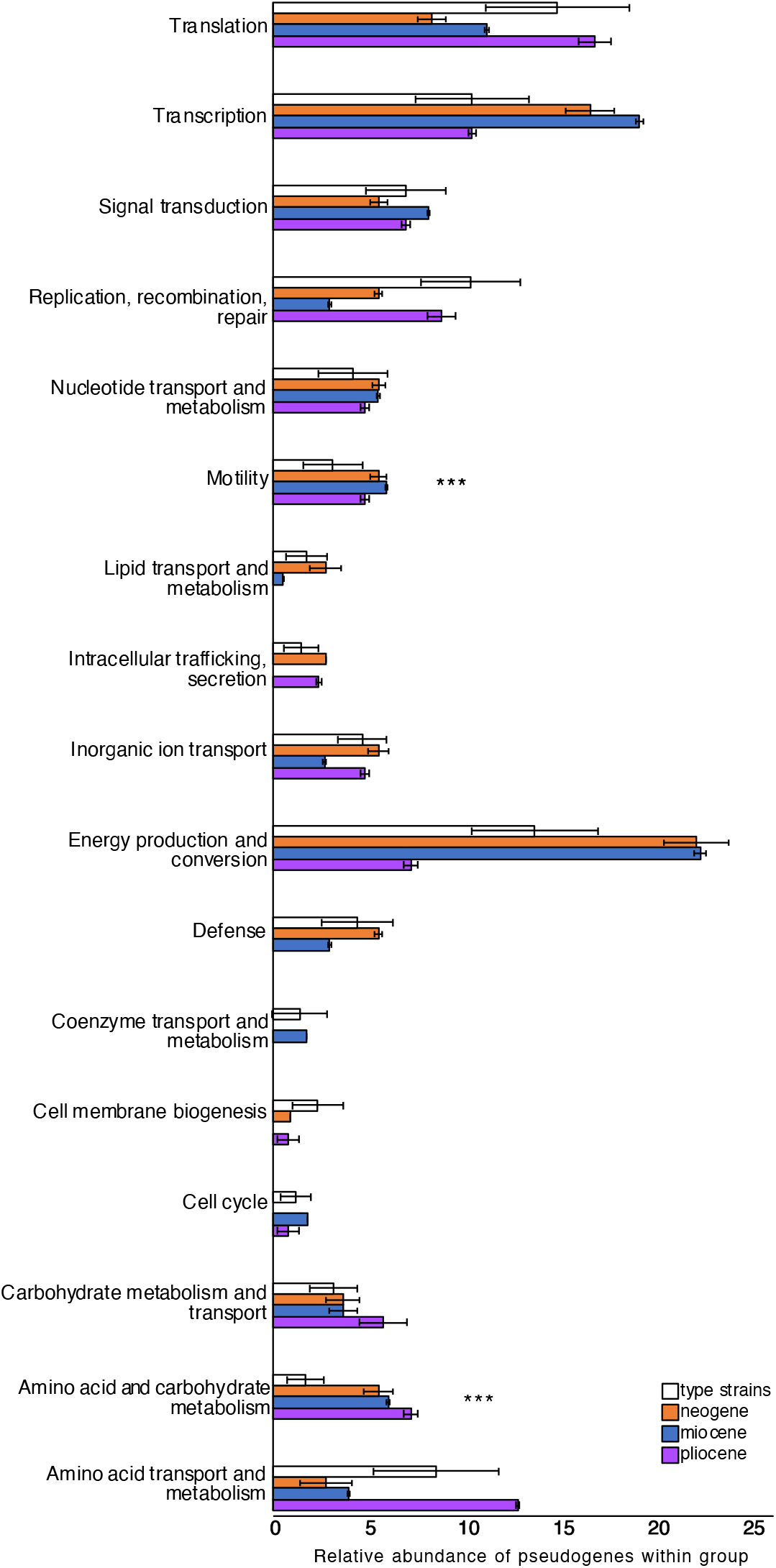
Histograms showing the average relative abundance of functional categories in pseudogenes found within each of the three subseafloor populations, compared to the type strains. The error bars represent standard deviations, and asterisks indicate functional categories of pseudogenes that were significantly higher in the subseafloor genomes compared to the type strain genomes (two sided T-Test, P<0.01).

## Materials and Methods

### Sampling and pore water chemistry

All samples were taken during Expedition KN223 of the *R/V Knorr* in the North Atlantic, from 26 October to 3 December 2014. At site 11 (22°47.0’ N, 56°31.0’ W, water depth ~5600 m) via a long core piston-coring device (~28 m). Additional details of sampling are published elsewhere (*11, 13*). Dissolved oxygen concentrations in the core sections were measured with optical O_2_ sensors from the equilibrated core sections and measured with needle-shaped optical O_2_ sensors (optodes) (PreSens, Regensburg, Germany) as described previously (*11, 13*). The dissolved O_2_ data from Expedition KN223 are archived and available online in the Integrated Earth Data Applications (IEDA) database (http://www.iedadata.org/doi?id=100519).

### Physical sediment properties

Deep-sea abyssal clay is characterized by very low permeability and extremely small pore diameter, despite its high porosity (*15*). Deep sea clay particles have a grain size of <0.2 μm, and the pore space between clay particles is smaller than a bacterial cell, limiting the movement of bacteria through pore space in the clay. Bioturbation can vertically redistribute cells within marine sediment, but bioturbation is restricted to the upper 0.5 meters of sediment (*31*), and thus cannot vertically transport sediment surface material and microbes to depths of 3 and 6 mbsf. Considering the mean sedimentation rate of 1 m per million years, it can be concluded that the bacterial cultures obtained from sediment collected at 3 and 6 mbsf have been physically isolated from the surface world for millions of years.

### DNA extraction, qPCR, 16S rRNA gene sequencing

DNA extractions, qPCR, and 16S rRNA gene sequencing were performed previously and described in Vuillemin et al (*12*). In brief, subcores were sampled aseptically with sterile syringes were subsampled aseptically in an ultraviolet (UV)–sterilized DNA/RNA clean HEPA-filtered laminar flow hood. DNA extraction was extracted from 10 g of sediment transferred into 50 ml of Lysing Matrix E tubes (MP Biomedicals) containing silica glass beads and homogenized for 40 s at 6 m/s using a FastPrep-24 5G homogenizer (MP Biomedicals) in the presence of 15 ml of preheated (65°C) sterile-filtered extraction buffer [76 volume % 1 M NaPO4 (pH 8), 15 volume % 200 proof ethanol, 8 volume % MoBio’s lysis buffer solution C1, and 1 volume % SDS]. The samples were incubated at 99°C for 2 min and frozen overnight at −20°C, thawed, and frozen again at −20°C overnight, followed by additional incubation at 99°C for 2 min and a second homogenization using the settings described above. After the second homogenization, the samples were centrifuged for 15 min, and the supernatants were concentrated to a volume of 100 ml using 50-kDa Amicon centrifugal filters (Millipore). Coextracted PCR-inhibiting humic acids and other compounds were removed from the concentrated extract using the PowerClean Pro DNA Cleanup Kit (MoBio). Extraction blanks were performed alongside the samples to assess laboratory contamination during the extraction process.

DNA was quantified fluorometrically using a Qubit with a double-stranded DNA high-sensitivity kit (Life Technologies). qPCR was performed using the custom primer dual indexed approach that targets the V4 hypervariable region of the 16S rRNA gene using updated 16S rRNA gene primers 515F/806R (515F, 5’-GTGYCAG-CMGCCGCGGTAA-3’; 806R, GGACTACNVGGGTWTCTAAT) (*32*). Barcoded V4 hypervariable regions of amplified 16S rRNA genes were sequenced on an Illumina MiniSeq following an established protocol (*33*). Bioinformatic processing of these previously published sequence data is described by Vuillemin et al (*12*) in detail.

### Long term incubation set up

Prior to setting up the incubations, the subcores were sampled with sterile syringes using the sample aseptic technique used for the DNA extraction. For each sample depth, seven grams of abyssal clay was placed into sterile 20-mL glass flasks and incubated with 4 mL of sterile artificial seawater composed of either H_2_^18^O (97% atomic enrichment) or unlabeled artificial seawater. Vials were crimp sealed, with an oxygenated headspace of approximately 10 mL, and incubated at 8 °C. The artificial seawater was different from the porewater at depth because there was no added nitrate, but there was also no added ammonia which should be similar to the *in situ* conditions where ammonia is generally below detection (*12*). Oxygen was measured continuously throughout the incubations using non-invasive fiberoptic measurements as described previously (*12*). Small fluctuations in the oxygen measurements in the killed control, and experimental incubations, were likely due to temperature fluctuations of the incubator itself (±1°C), since the non-invasive fiber optic oxygen sensor spots are temperature sensitive (*12*). Oxygen consumption was detectable over 18 months in slurries consisting of sediment and sterile artificial seawater (Fig S1), suggesting the presence of actively respiring microbes.

We used qSIP to measure the atom % ^18^O-enrichment of actively growing microbial taxa as described previously (*12*). In brief, after 7 and 18 months incubations DNA was extracted and subjected to Cesium Chloride (CsCl) density gradient centrifugation. The same 16S 515F/806R primers (described above) were used in qPCR (described above) to determine density shifts in the peak DNA of buoyant density (BD) for each incubation. 16S rRNA gene amplicons from each fraction resulting from the density gradient fractionation were Illumina sequenced as described previously (*12*). To identify contaminants that may have entered during the fractionation process, we also included in the sequencing run extraction blanks from the SIP fractionation. OTUs containing sequences from extraction blanks were removed. Excess atm% ^i8^O-enrichment was calculated for each OTU (including OTU6, corresponding to the subseafloor *Thalassospira*) according to the equations for quantifying per OTU atomic enrichment.

The number of doublings for the *Thalassospira* OTU (OTU_6) detected at the 18 month timepoint was calculated using qPCR normalized relative abundance of the 16S rRNA genes at T0 and 18 months. The number of doublings was divided by the total number of days incubated to calculate doubling times in days.

### Enrichments, cultivation, and sub-cultivation

After the 18 months of incubation in sterile ^18^O-labeled artificial seawater, 25 μL of slurry was plated onto solid media (10 mg/mL yeast extract and 8 mg/mL agar in artificial seawater [30 mM MgCl_2_ · 6H_2_O, 16 mM MgSO_4_ · 7H_2_O, 2 mM NaCO_3_, 10 mM KCl, 9 mM CaCl_2_, 450 mM NaCl)]), and after 2 days incubated in the dark at room temperature, abundant colonies were observed growing on the surface of the petri dishes (Fig S1). No colonies were observed to grow on control petri dishes that received 25 μL of ^18^O-labeled artificial seawater slurry incubated for 18 months using starting material from autoclaved sediment (killed controls). This indicated that the colony-forming bacteria were from the sediment and not contaminants introduced during the experimental set up of the incubations. We attempted to culture chemoheterotrophic microbes directly from the collected sediment samples using the same conditions, but no colony-forming units were observed on the petri dishes, even after several months of incubation. Thus, long-term incubation of the sediment at 8 °C simply in the presence of added water apparently stimulated the activity of many subseafloor bacteria to a point at which they were able to grow on the surface of a petri dish.

Twelve colonies were picked from petri dishes containing colonies from the 3-mbsf and 6-mbsf slurries. These colonies were streaked individually onto twelve separated new petri dishes, and a single colony was picked from each of the twelve petri dishes (representing an original colony forming unit from the enrichment) and grown in sterile liquid media (10 mg/mL yeast extract and 8 mg/mL agar in artificial seawater [30 mM MgCl_2_ · 6H_2_O, 16 mM MgSO_4_ · 7H_2_O, 2 mM NaCO_3_, 10 mM KCl, 9 mM CaCl_2_, 450 mM NaCl)]). Single colonies were then grown up in liquid media. A portion of each of these colonies was used for DNA extraction and genome sequencing, and the remaining volume was frozen as glycerol stocks. Growth rates were determined in experiments with 20 mL crimp sealed glass flasks containing 0.1 mL of glycerol stock inoculated into 10 mL liquid media (10 mg/mL yeast extract in artificial seawater [30 mM MgCl_2_ · 6H_2_O, 16 mM MgSO_4_ · 7H_2_O, 2 mM NaCO_3_, 10 mM KCl, 9 mM CaCl_2_, 450 mM NaCl)]) with 10 mL headspace, and gentle shaking. The optical density at 600 nm was measured once every 30 minutes with a spectrophotometer, and growth rates were calculated from the exponential phase. Under these conditions, growth rates of the subseafloor *Thalassospira* were similar across all three clades and ranged from 0.064 to 0.31 h^−1^ (Fig. S5).

### Assessing the possibility for genome evolution during the 18 month enrichment

Because bacteria can evolve on lab experimental timescales (*5, 6*) we considered the possibility that all diversification and evolution happened during the 1.5 year enrichment. Using the qPCR-based estimate for doubling time of the subseafloor *Thalassospira* OTU (OTU_6) in the incubation which was 36 (± 1.5) days, the number of doublings with this rate over this time period would be approximately 15. According to “Drake’s rule (*20*), bacteria experience on average one mutation per 300 genomes replicated; thus, the amount of nucleotide diversity (hundreds to thousands of mutations: Fig S8) that could be accumulated during the incubation is insufficient to explain the observed divergence between the three subseafloor populations. We thus conclude that the inter-population nucleotide diversity resulted from mutations that were acquired after they were buried.

### Genome sequencing, de novo assembly, and annotation

DNA was extracted from the isolates grown in liquid culture until the end of exponential phase as described above. After reaching stationary phase, cultures were pelleted via centrifugation and the supernatant was decanted. The cell pellets were resuspended in a preheated (65°C) sterile filtered extraction buffer [76 volume % 1 M NaPO4 (pH 8), 15 volume %200 proof ethanol, 8 volume %MoBio lysis buffer solution C1,and 1 volume % SDS], and added to lysing matrix E tubes (MP Biomedicals) containing silica glass beads and homogenized for 40 s at 6 m/s using a FastPrep-24 5G homogenizer (MPBiomedicals). The samples were centrifuged for 15 min, and the dissolved high molecular weight DNA in the supernatant was concentrated to a volume of 100 μL using 50-kDa Amicon centrifugal filters (Millipore). The concentrated extract was cleaned of proteins and other non-genomic DNA organic matter using the PowerClean Pro DNA Cleanup Kit (MoBio). Extraction blanks were performed alongside the samples to assess laboratory contamination during the extraction process. Genomic libraries were prepared using the Nextera XT DNA Library Prep Kit (Illumina). Quality control and quantification of the libraries were obtained on an Agilent 2100 Bioanalyzer System using the High Sensitivity DNA reagents and DNA chips (Agilent Genomics). Metagenomic libraries were diluted to 1 nM using the Select-a-Size DNA Clean and Concentrator MagBead Kit (Zymo Research) and pooled for further sequencing on the Illumina MiniSeq platform. Genomic libraries were sequenced to a depth of ca. 100x coverage using a high-output paired end 2 x 150 sequencing regent kit (Illumina).

In addition to Illumina sequencing, the high molecular weight genomic DNA was sequenced using the NanoPore MinION. Sequencing libraries for the MinION were prepared using the Ligation Sequencing kit (Oxford NanoPore Technologies), according to the manufacturers instructions. Barcoded libraries were sequenced on the MinION using a Flongle R9 flow cell, base-called and demultiplexed using the MinIT with ont-minit-release v19.12.5 and ont-guppy-for-minit v3.2.10 for base calling (Oxford NanoPore Technologies).

A hybrid assembly was performed using both the short (Illumina) and long (NanoPore) read sequencing data using Unicycler (v.0.4.0), which uses *de novo* assembled Illumina data from SPADES to polish the *de novo* assembled contigs obtained from NanoPore data using RACON (*34*). The combined assemblies of Illumina and NanoPore data resulted in a relative low number of contigs (9-12 per genome), and a predicted genome completeness of 100% of nearly all genomes (Table S1). Genome completeness was determined using CheckM (*35*). Genomes were annotated using RASTk (*36*).

### Core genome phylogenetic analyses

The core genome was defined as the set of orthologous genes which were shared in all subseafloor and extant *Thalassospira* genomes. Orthologous genes were defined as those sharing >30% amino acid similarity to the collective suite of genes encoded within the type strain *Thalassospira xiamenensis* M-5. *T. xiamenensis* M-5 was chosen as the reference genome for this purpose, because it is the only publicly available genome of a cultivated *Thalassospira* isolate that is completely closed and represents a single chromosome and a 190 Kb plasmid (*14*). A total of 1,809 orthologous genes were identified that are encoded by all *Thalassospira* strains that had >30% sequence similarity to genes encoded within the *T. xiamenensis* M-5 genome. Each of these 1,809 genes was individually aligned between all *Thalassospira* strains using MUSCLE (*37*), and the individual 1,809 alignments were then concatenated into a single core genome alignment for the subsequent phylogenomic analysis (ClonalFrameML, HyPhy, aBRSEL) using Geneious Prime (version 2019.2.1). After concatenation of all core genes, the total size of the core genome alignment was 1,817,073 nucleotide characters, and 34 taxa (21 subseafloor strains, and 13 type strain taxa). A Maximum-Likelihood phylogeny was created using PhyML (*38*) with a GTR model of evolution and 100 bootstrap replicates, which was implemented within SeaView (*39*). The resulting phylogenetic tree and the concatenated core genome alignment were used as inputs for subsequent ClonalFrameML and dN/dS analyses.

The contributions of mutations and recombination to the genomic diversity in the concatenated core genome alignment, the number of recombination events (imports) per genome, and the positions of recombination hot spots, were investigated using ClonalFrameML (*19*). Nucleotides unaffected by recombination are referred to as unimported and nucleotides subject to recombination are referred to as imported (*19*). ClonalFrameML provides the relative rate of recombination to mutation (R/Theta), the mean length of recombined DNA (Delta), and the mean divergence of imported DNA (Nu). These results were used to calculate the relative contribution of recombination versus mutation to the overall genomic diversity (r/m), using the formula r/m = (R/Theta) * Delta * Nu. ClonalFrameML was performed in three separate runs, respectively containing a core genome alignment that contained (1) all genomes, (2) only the subseafloor genomes, and (3) only the type strains. The resulting r/m values from these three groups (presented in Table 1) were then used to interpret the relative importance of mutations compared to recombination, in the separate groups (e.g., type strains versus subseafloor strains).

In addition to calculating sites and rates of recombination in the core genome, ClonalFrameML also estimates the ancestral sequences at internal nodes of the clonal genealogy, and any missing base calls in the observed sequences. The reconstruction of ancestral sequence states is performed using maximum likelihood and the ClonalFrame model can be thought of as a hidden Markov model (HMM) when the ancestral and descendant genomes for each branch of the clonal genealogy have been observed or reconstructed (*19*). The hidden state of the HMM records whether each nucleotide was subject to recombination or not on the branch connecting the two genomes. We acknowledge that drawing inference under the resulting ancestral recombination graph is a notoriously complex statistical problem (*19*). Instead, here we use ClonalFrameML only to assess within-group recombination (e.g., between species within the genus *Thalassospira*), and thus our analysis cannot assess the influence of external recombination (from species outside the genus *Thalassospira*).

The ratio of non-synonymous (dN) to synonymous (dS) mutations in the core genome alignment (global ω ratio) was estimated using HyPhy v2.2.4 (*40*), and applying the adaptive branch-site random-effects likelihood (aBSREL) approach (*41*) to all branches in all subfamilies. Because of the high similarity of the subseafloor genomes, aBSREL was run multiple times using the core genome alignment with only one representative of the nearly identical subseafloor genomes included in each separate run. For each of these runs, one representative genome of *T. xiamenensis* strain ‘Neogene’, *Txiamenensis* strain ‘Miocene, and ‘Ca. *T. pliocenensis*’ were included together with all other *Thalassospira* type strain genomes in each aBSREL run. Then, the aBSREL run was repeated with the same type strains but different subseafloor genomes from those same three clades, until dN/dS estimates were obtained for all subseafloor genomes.

### Pangenome analysis

All subseafloor and extant *Thalassospira* genomes were analyzed in Anvi’o v6.2 using pangenome workflow (*42*). Briefly, each genome was converted into an anvi’o contigs database. Genes were functionally annotated using eggnog v5.0 (*43*) with eggNOG-mapper (*44*) and imported back to each genome’s anvi’o contig database. Genome storages were generated using ‘anvi-gen-genomes-storages’ and ‘anvi-pan-genome’ was deployed with parameters ‘--min-bit 0.5’ (*45*), ‘--mcl-inflation’ 10 (*46*), and the flag ‘--use-ncbi-blast’ (*47*). The anvi’o pan database and summary of gene clusters stored in FIGshare (https://figshare.com/s/06ba1287a00ab01a1ee).

### Identifying pseudogenes

We estimated the number of pseudogenes within the genomes using two programs, Psi-Phi (*48*) and DFAST (*49*). Psi-Phi uses a conservative criterion considering a pseudogene only when it lost >20% of its original length, and enhances pseudogene recognition among closely related strains both in annotated regions by identifying incorrectly annotated open reading frames (ORFs) and in intergenic regions by detecting new pseudogenes (*48*). Psi-Phi classifies pseudogenes as either identified pseudogenes and those as being possible, but potentially not pseudogenes. To be conservative, we only considered genes identified as pseudogenes from Psi-Phi and did not consider those flagged as ‘potential pseudogenes’. As a second check of pseudogene content, we searched genomes for pseudogenes using DFAST (*49*). The estimated number of pseudogenes per genome was then taken as an average of the numbers detected both using both methods (Psi-Phi and DFAST). On average, Psi-Phi identified a higher number of pseudogenes per genome (57 ±10) compared to DFAST (32 ±4), but the variation between methods for the same genome was consistent (average variation =27, standard deviation of averages = 7). This minimal variation between individual genomes indicates that biases inherent to the pseudogene prediction methods affected the different genomes equally, and thus allow for a pseudogene comparison between the genomes.

